# *RB1* deletion in RB-pathway disrupted cells results in DNA damage and cancer progression

**DOI:** 10.1101/564567

**Authors:** Aren E. Marshall, Michael V. Roes, Daniel T. Passos, Megan C. DeWeerd, Andrea C. Chaikovsky, Julien Sage, Christopher J. Howlett, Frederick A. Dick

## Abstract

Proliferative control in cancer cells is frequently disrupted by mutations in the RB-pathway. Intriguingly, *RB1* mutations can arise late in tumorigenesis in cancer cells whose RB-pathway is already compromised by another mutation. In this study, we present evidence for increased DNA damage and instability in *CDKN2A* silenced cancer cells when *RB1* mutations are induced. We generated isogenic *RB1* mutant genotypes with CRISPR in a number of cell lines. Cells with even one mutant copy of *RB1* have increased basal levels of DNA damage and increased mitotic errors. Elevated levels of reactive oxygen species as well as impaired homologous recombination repair underlie this DNA damage. When xenografted into immune compromised mice *RB1* mutant cells exhibit an elevated propensity to seed new tumors in recipient lungs. This study offers evidence that late arising *RB1* mutations can facilitate genome instability and cancer progression that are beyond the pre-existing proliferative control deficit.

## Introduction

Loss of proliferative control is a defining feature of human cancer. Most cancer cells develop cell intrinsic mechanisms of supplying growth stimulatory signals as well as disrupting the response to cell cycle arrest cues (Hanahan and Weinberg, 2011). To this end, mutations in the RB-pathway are central to disrupting proliferative control in tumorigenesis (Burkhart and Sage, 2008; Knudsen and Knudsen, 2008; Sherr and McCormick, 2002). Deletion of the *RB1* gene prevents cell cycle arrest in response to a broad range of signals (Knudsen and Knudsen, 2008). Similarly, overexpression or hyperactivation of D-type Cyclins and their associated Cyclin Dependent Kinases (CDKs) can lead to constitutive RB protein (RB) phosphorylation and cell cycle entry. Lastly, deletion or promoter methylation of *CDKN2A* that encodes the CDK inhibitor p16 serves to deregulate kinase activity causing constitutive phosphorylation of RB. Cancer cell genomes that sustain a single mutation in this pathway are considered to have disrupted RB-pathway function and are deficient for cell cycle control (Dyson, 2016; Knudsen and Knudsen, 2008; Sherr, 1996). Historically, this concept of RB-pathway inactivation suggested that mutations in different components of the pathway are relatively equivalent and additional mutations provide no subsequent advantage to cancer progression (Dick et al., 2018; Knudsen and Knudsen, 2008; Sherr, 1996; Sherr and McCormick, 2002).

A number of recent clinical observations challenge the logic of single RB-pathway mutations in cancer. First, multiple studies have shown that *RB1* loss is specifically predictive of a favourable response to chemotherapy (Cecchini et al., 2015; Garsed et al., 2017; Ludovini et al., 2004; Zhao et al., 2012), whereas p16 expression or overall proliferative rates are not (Cecchini et al., 2015; Garsed et al., 2017; Zhao et al., 2012). This suggests that RB-pathway mutations are not necessarily equivalent. Second, a number of studies have suggested that *RB1* gene loss is more prevalent in advanced cancers, or mechanistically contribute to progression or dissemination (Beltran et al., 2016; McNair et al., 2017; Robinson et al., 2017; Thangavel et al., 2017), a stage where cell autonomous proliferative control is presumably already deregulated. Collectively, these examples suggest *RB1* mutation facilitates more than alterations to proliferative control and that *RB1* loss may confer other cancer relevant characteristics. Remarkably, some studies even highlight that single copy loss of *RB1* may be functionally significant (Coschi et al., 2014; Gonzalez-Vasconcellos et al., 2013; McNair et al., 2017; Zheng et al., 2002).

Beyond the RB protein’s role in cell cycle control through E2F transcriptional regulation, it has been reported to participate in a host of functions that contribute to genome stability (Velez-Cruz and Johnson, 2017). These include chromosome condensation through RB-dependent recruitment of Condensin II and Cohesin (Longworth et al., 2008; Manning et al., 2014). The RB protein also influences repair of DNA breaks through both non-homologous end joining (NHEJ) (Cook et al., 2015), and homologous recombination (HR) (Velez-Cruz et al., 2016), and induction of mitochondrial biogenesis that impacts cell metabolism (Benevolenskaya and Frolov, 2015; Jones et al., 2016; Nicolay et al., 2015). Some of these functions, such as repair of DNA breaks by HR, are obligatorily outside of RB’s role in G1 to S-phase regulation.

In addition, other roles, such as effects on mitochondrial biogenesis and metabolism take place in proliferating populations of cells further suggesting that this is independent of G1-S regulation and the RB-pathway. It is noteworthy that some atypical RB functions in genome stability, or late stage cancer progression, may be sensitive to single copy loss (Coschi et al., 2014; Gonzalez-Vasconcellos et al., 2013; Sharma et al., 2010; Zheng et al., 2002). Thus, the existence of shallow *RB1* deletions may indicate that RB’s less well appreciated functions in genome stability could underlie cancer relevant characteristics that are independent of classical RB-pathway function in cancer (Dick et al., 2018).

In order to test if *RB1* loss is relevant to cancer cells that already possess RB-pathway disruption, we induced mutations in *RB1* using CRISPR/Cas9. These cells displayed spontaneous DNA damage as evidenced by γH2AX foci and elevated levels of reactive oxygen species. We also determined that *RB1* mutations decreased the ability to repair DNA breaks by homologous recombination, and this is supported by elevated levels of anaphase bridges in mitosis. *RB1* mutant cells were xenografted into immune compromised mice and this revealed similar growth kinetics in subcutaneous implantation, with *RB1* null showing greater propensity to colonize lungs. These experiments underscore the discovery that *RB1* mutation in cells that already possess RB-pathway disruption creates DNA damage and fuels cancer progression.

## Results

### Spontaneous DNA damage in *RB1* deficient cancer cells

To investigate *RB1* deficiency in RB-pathway disrupted cells, we utilized U2OS cells that do not express p16, the product of *CDKN2A* (Forbes et al., 2017). We used CRISPR technology and gRNA pairs that target exon 22 of *RB1* because loss of this exon creates null alleles in cancer (Horowitz et al., 1990)(SFig. 1A). Cells were transfected with plasmids to deliver pairs of gRNAs and Cas9 (or the D10A mutant). Following transient drug selection, colonies were isolated, expanded, and genotyped by PCR to search for *RB1* deletions (SFig. 1B). Candidates were rigorously selected by checking RB protein expression (SFig. 1C), ensuring heterozygous clones were not cell mixtures (SFig. 1D), and confirming that the most likely off targets were not mutated (SFig. 1E). Using this approach, we selected four clones each for wild type and knockout *RB1* genotypes, and three clones for the *RB1*^+/−^ genotype that were used in subsequent experiments.

To determine if *RB1* mutation status affects genome stability in these engineered cell lines, DNA damage was assessed in untreated, proliferating cells by staining for γH2AX. Foci were visualized by immunofluorescence microscopy and images were captured using confocal microscopy (Fig. 1A). The quantity of foci per nucleus was determined and this revealed a significant increase in γH2AX in the knockout and heterozygous lines compared to those that are wild type for *RB1* (Fig. 1B). To extend this observation, we analyzed γH2AX foci in control and *RB1* null U2OS clones produced by targeting a different exon 2, as well as clones of control and *RB1* mutant NCI-H460 *(CDKN2A* deleted) and NCI-H1792 *(CDK4* amplified) lung cancer cells (SFig. 2). This analysis also demonstrated an elevation in the number of γH2AX foci in each of the *RB1* deleted lines.

**Figure 1.**
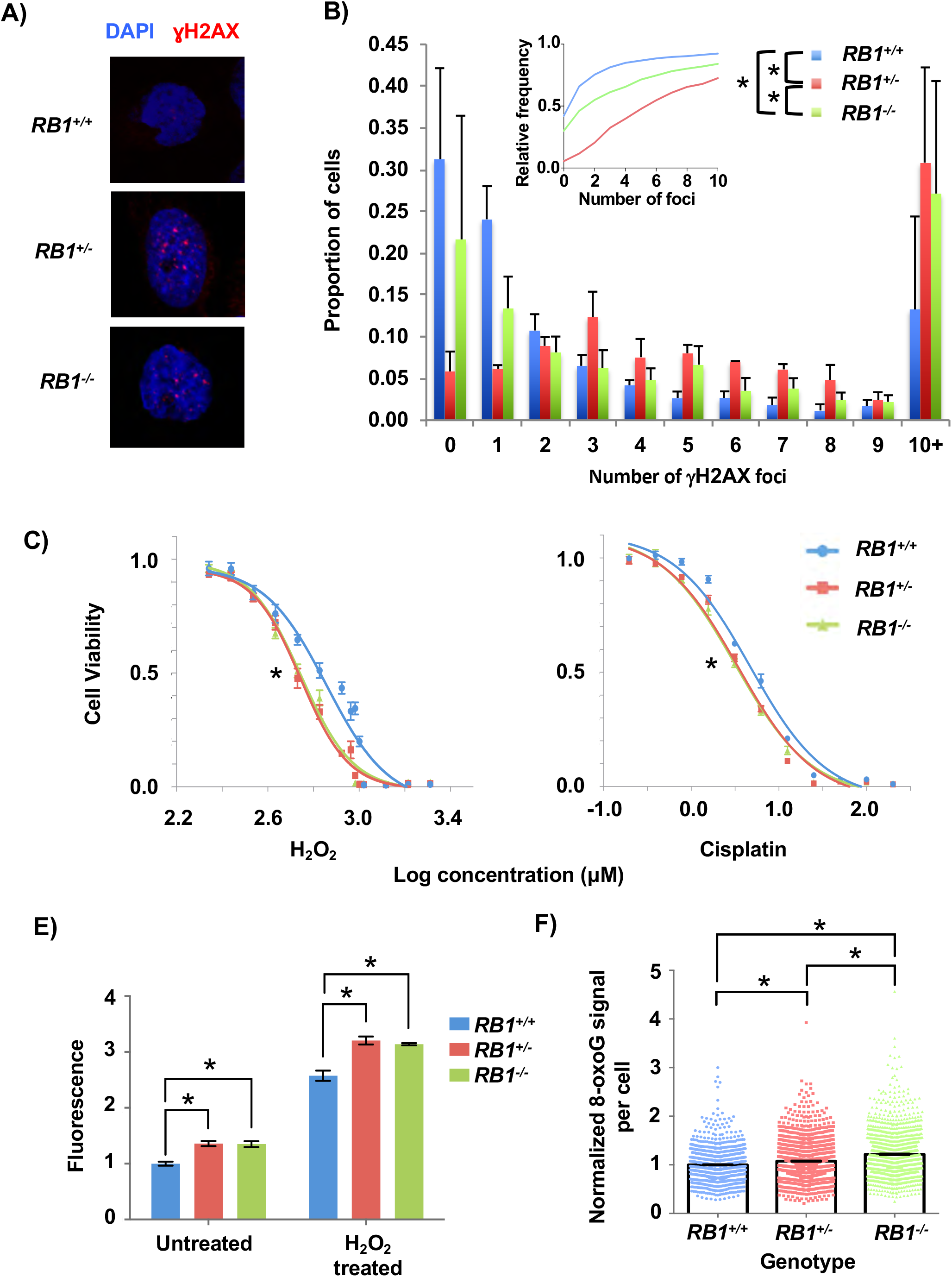
U2OS cells deficient for one copy of *RB1* have increased DNA damage, elevated reactive oxygen species, and sensitivity to cisplatin. **(A)** Representative confocal microscopy images of γH2AX foci (red) in control, heterozygous, and homozygous *RB1* mutant cells. Cells were counterstained with DAPI to visualize nuclei (blue). **(B)** γH2AX foci counts for each of the *RB1* genotypes. The average proportion of cells with discrete numbers of foci are shown as histograms, while the cumulative frequency of foci for each genotype is shown as an inset. The distribution in foci for *RB1* wild type (4 different clones), heterozygous (3 different clones) and knockouts (4 different clones) are significantly different as determined by the Kolmogorov-Smirnov test. **(C)** Hydrogen peroxide and **(D)** cisplatin were added to cultures of the indicated genotypes of cells. Viability was assessed after 72 h using alamarBlue and dose response curves were used to calculate IC50 values for each genotype. Both *RB1* mutant genotypes are significantly lower than U2OS cells (as determined by one-way ANOVA). **(E)** To detect reactive oxygen species (ROS), CA-DCF-DA was added to culture media at the end of 72 h of mock treatment or hydrogen peroxide. Normalized fluorescence was averaged for four clones of *RB1* wild type and knockout genotypes, and three clones for the heterozygous genotype. Mean values were compared by two-way ANOVA. **(F)** Cells were fixed and stained for 8-oxoG and DAPI and visualized by fluorescence microscopy. The average 8-oxoG signal per nucleus was determined using ImageJ with DAPI staining defining nuclear area. Three clones per genotype were used and data was normalized to the mean signal from *RB1* wild type. Statistical significance in staining intensity was determined by Kruskal-Wallis one-way analysis of variance and Dunn’s multiple comparisons test. All error bars are ±1 SEM. *\*P* < 0.05.

To further investigate the source of DNA damage in *RB1* mutant cells, we assessed their sensitivity to a number of chemical agents to determine if specific stresses could amplify defects that cause increased DNA damage. We tested aphidicolin, a DNA polymerase inhibitor, that causes replication stress and etoposide, a topoisomerase inhibitor, that creates DNA double stranded breaks. Hydrogen peroxide (H_2_O_2_) was used to induce oxidative damage, and cisplatin was used to create interstrand cross links, among other damaging effects. Representative clones of each genotype were treated for 72 h, after which alamarBlue was used to quantitate the cytotoxicity of each agent. These assays revealed that both heterozygous and homozygous *RB1* mutations sensitize cells to hydrogen peroxide and cisplatin (Fig. 1C&D), but not aphidicolin or etoposide (SFig. 3A&B). This suggests oxidative damage may underlie some aspects of the DNA damage phenotype in U2OS cells. We compared ROS levels in wild type and *RB1* mutant cells with and without H_2_O_2_ using a ROS indicator, 5(6)-carboxy-2’,7’-dichlorodihydrofluorescein diacetate (CA-DCF-DA). For both untreated and H_2_O_2_ treated cells, there was more fluorescence of the ROS indicator in *RB1* mutant cells, as *RB1*^−/−^ and *RB1*^+/−^ were equivalent (Fig. 1E). We also fixed and stained cells for 8-oxoguanine (8-oxoG), one of the most abundant lesions resulting from oxidative modification of DNA (Furtado et al., 2012), and quantified the staining in DAPI-defined nuclear area using ImageJ (Schneider et al., 2012). Control U2OS values were used to normalize the 8-oxoG signal from *RB1* mutants. Again, both *RB1*^+/−^ and *RB1*^−/−^ U2OS cells had more 8-oxoG staining than the *RB1*^+/+^ cells (Fig. 1F).

These experiments indicate that loss of *RB1* in cells with pre-existing RB-pathway defects increases basal levels of DNA damage. Reactive oxygen species appear to be one source of this damage. Chemical agents that directly induce breaks did not selectively affect *RB1*mutant cells, which was consistent with a lack of elevation in 53BP1 foci in *RB1*-mutant U2OS cells compared to controls (SFig. 3C). This suggests the nature of DNA damage marked by γH2AX in these *RB1*-mutant cells is not necessarily double stranded DNA breaks, or double stranded DNA breaks are not repaired by NHEJ. Overall, these experiments indicate that *RB1* loss contributes to an unstable genome, regardless of the proliferative control status of the cell.

### *RB1* mutant cells have randomly distributed DNA damage

To further understand spontaneous DNA damage in *RB1* mutant U2OS cells, we sought to determine if damage occurred at specific locations within the genome. We performed ChIP-sequencing to identify DNA sequences associated with γH2AX, as well as histone H4 as a control. Because spontaneous damage in untreated cell cultures is relatively inabundant, we pooled chromatin from 20 separate γH2AX ChIP experiments per genotype to create each sequencing library (Fig. 2A-E). We determined peak locations and number using MACS (Zhang et al., 2008), and the quantity of γH2AX and H4 peaks were similar between genotypes (Fig. 2A). Looking at a large region of chromosome 4 as a representative view of the genome, we did not observe consequential differences between genotypes for γH2AX or H4 peaks (Fig. 2B). Since peak-finding at a genome scale failed to indicate obvious locations of DNA damage enrichment, we investigated individual genome sequence categories in search of elevated γH2AX deposition. The proportion of aligned γH2AX ChIP-Seq reads per million mapped reads versus input reads for each of the genotypes were compared within each repetitive element category and log2 transformed for display as a heat map (Fig 2C). Some categories, such as SINEs and multicopy genes appear to have slight enrichment of γH2AX localization in *RB1*^+/−^ and *RB1*^−/−^ compared to *RB1*^+/+^ based on color, but a two-tailed one-sample *t*-test with FDR multi-test correction did not score these as significant. Therefore, even within repetitive sequences in the genome, there does not seem to be an enrichment of γH2AX within *RB1* mutant cells compared to control.

**Figure 2.**
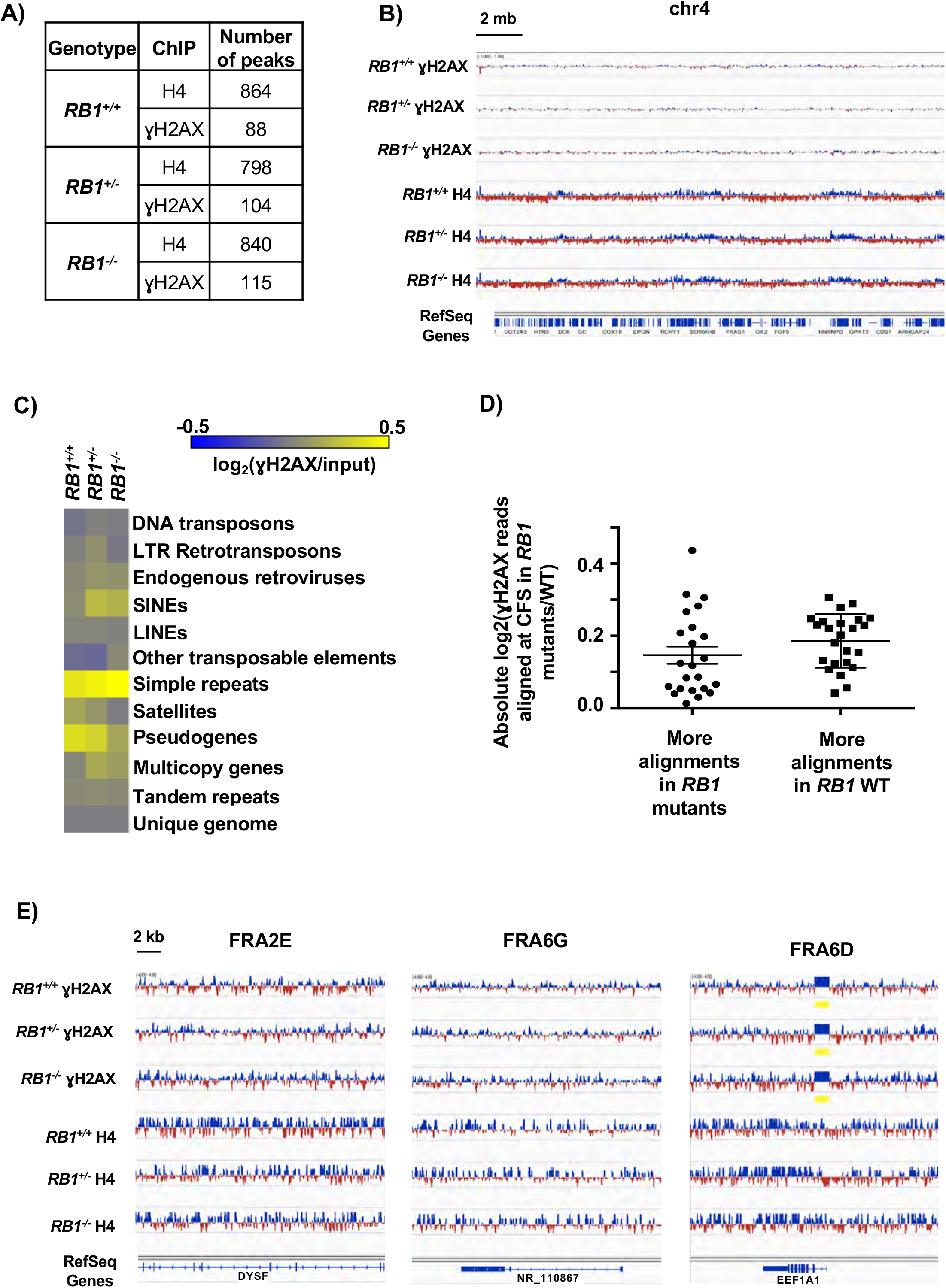
γH2AX is randomly distributed in the genomes of *RB1* mutant cells. **(A)** Total number of MACS peaks found for H4 control and γH2AX ChlP-Seq reads for the indicated genotypes. **(B)** A 20 Mb region of chromosome 4 is shown with ChIP-Seq read alignments for γH2AX and H4. Tracks were normalized by subtracting input reads. Blue indicates more reads in the ChIP versus input, red is the opposite. **(C)** The number of ChIP-Seq reads mapping to repetitive sequences, as well as unique genome regions, was determined. The heatmap shows the log2 ratios of the abundance of γH2AX precipitable reads per million mapped reads versus input for each of the respective genotypes at each element analyzed. **(D)** Aligned γH2AX ChIP-Seq read proportions within common fragile sites (CFS) were first normalized to their respective inputs, and then *RB1*^+/−^ and *RB1*^−/−^ were normalized to wild type and log2 transformed. A two-tailed one-sample *t*-test was performed to determine if the normalized mean read count proportions of the *RB1* mutants at the various CFS is equal to the normalized read count proportion of the corresponding CFS in the wild type. CFS where FDR were less than 0.1 were grouped according to whether there were significantly more alignments in the *RB1* mutants, or significantly more alignments in the *RB1*^+/+^ sample. There is no significant difference between these two categories (determined by unpaired t-test). Error bars are ±1 SEM. **(E)** ChIP-Seq tracks for γH2AX and H4 at representative CFS are displayed. FRA2E had more reads in *RB1*^+/−^ and *RB1*^−/−^ than the wild type, FRA6G had more reads in the wild type than the mutants, and FRA6D had no change in the proportion of γH2AX reads that aligned between the genotypes. Regions of significant enrichment (MACS peaks) are denoted by yellow bars.

Previous studies suggest that γH2AX levels can be elevated at common fragile sites (CFS) under standard cell culture conditions (Harrigan et al., 2011). To investigate these locations we quantified the number of γH2AX ChIP-Seq reads and scaled them to the proportions of total aligned reads and normalized them to input levels. *RB1*^+/−^ and *RB1*^−/−^ were compared to wild type using a two-tailed one-sample *t*-test. This analysis revealed 23 CFS that had more γH2AX alignments than the wild type and 24 CFS that had significantly less in *RB1*-mutant cells compared to controls (SFig. 4 and Fig. 2D). Fig. 2E shows examples of CFS locations with increased γH2AX (FRA2E) in mutants, reduced γH2AX (FRA6G) in mutants, and unchanged γH2AX levels (FRA6D). These examples appear highly similar between genotypes. Overall, it is possible that the distribution of γH2AX within each CFS may be shifting slightly between the mutants and the wild type. However, there does not appear to be more of a bias in general for γH2AX elevation at CFS in the *RB1* mutant cells.

Collectively, our analysis of γH2AX distribution across the genome suggests there is no particular chromosome location or sequence category that is preferentially enriched for this mark of DNA damage. These data suggest that the increase in γH2AX foci observed in *RB1*^+/−^ and *RB1*^−/−^ cells is likely due to an overall increase in DNA damage, and not newly arising locations, or “hotspots” of damage. This is in contrast to primary cells with a normal RB-pathway that experience RB loss and preferentially damage centromeric repeats (Coschi et al., 2014). The elevated sensitivity to peroxide and cisplatin, and increased ROS and 8-oxoG are consistent with DNA damage being randomly located in *RB1* mutant U2OS cells.

### Homologous recombination repair defects in *RB1* deficient cancer cells

Another potential source of intrinsic DNA damage could arise from defective repair. For this reason we investigated the efficiency of HR and NHEJ repair using fluorescent reporters (Bennardo et al., 2008; Pierce et al., 1999). In this assay a promoterless, but functional, GFP is used to repair an adjacent break induced in a mutant, expressed, form of the GFP gene (Fig. 3A). We generated stable U2OS lines bearing this reporter and deleted *RB1* with lentiviral delivery of Cas9 and an *RB1* specific gRNA (Fig. 3B). Introduction of the restriction enzyme I-SceI into these cells induced breaks and *RB1* deficient cells were defective for their repair (Fig. 3C). Similarly, we generated U2OS cells that stably maintain an NHEJ reporter for repair of an induced break that links a constitutive promoter with a GFP gene. Loss of RB expression was again confirmed by western blotting (Fig. 3E), and induction of breaks was used to test repair in an *RB1* deficient background (Fig. 3F). This failed to reveal a defect in repair suggesting that *RB1* loss in U2OS cells specifically reduces HR repair.

**Figure 3.**
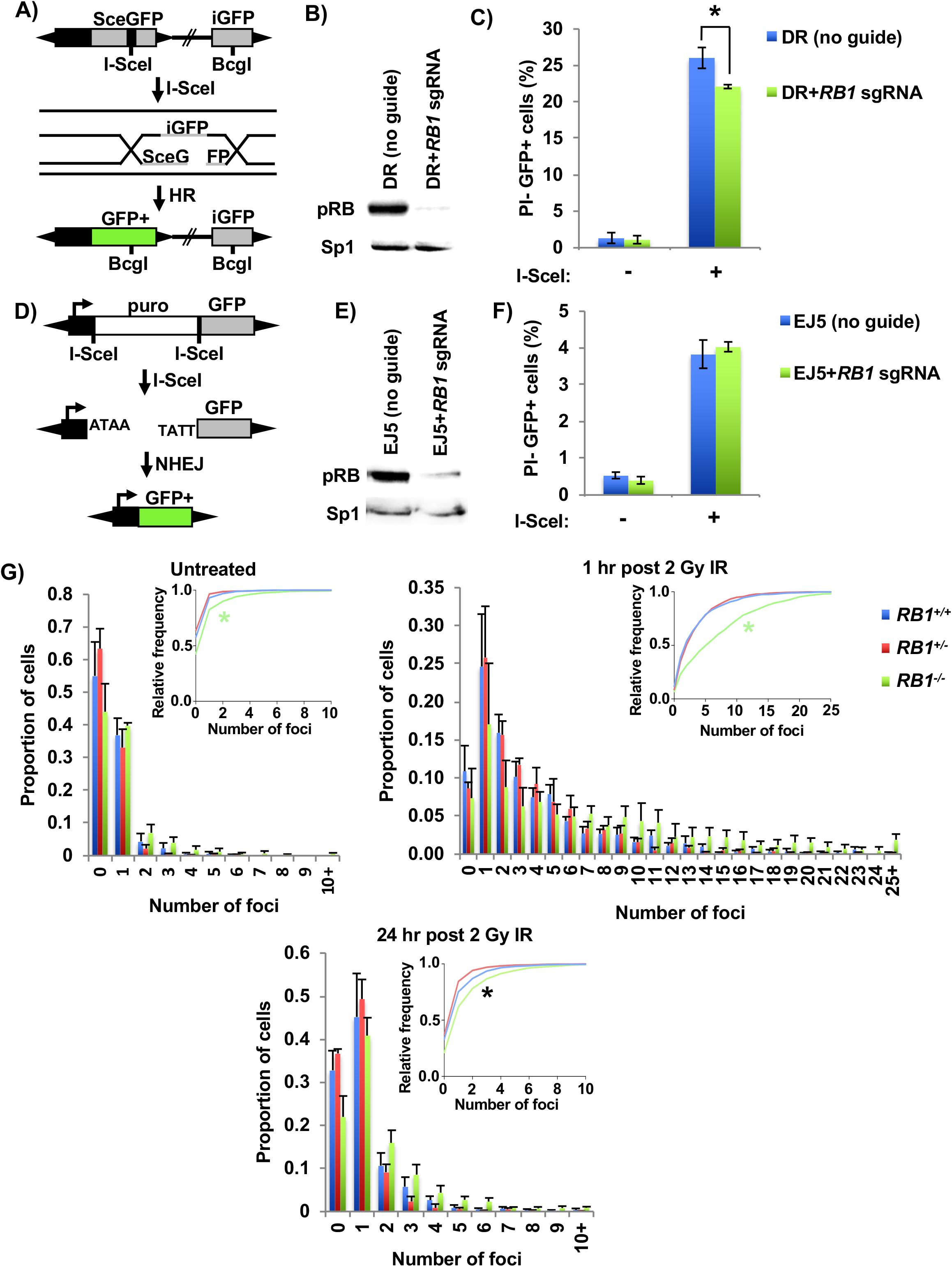
Defective homology directed repair of breaks in *RB1* mutant cells. **(A)** Schematic of the DR homology directed repair construct used. Cleavage of an I-SceI site integrated into an expressed, but mutant GFP gene, can be repaired from a downstream internal GFP fragment (iGFP). **(B)** U2OS cells with clonal integration of the HR reporter construct, were ablated for *RB1* expression with lentiviral delivery of Cas9 and an *RB1* specific sgRNA. Relative expression of RB was determined by western blotting and SP1 serves as a loading control. **(C)** HR repair efficiency of *RB1* mutant U2OS cells was determined by transfecting an expression vector for I-SceI endonuclease (I-SceI +), or relevant negative control expression vector, and quantitating PI- and GFP+ cells by flow cytometry (n=5). **(D)** Schematic of the EJ5 NHEJ reporter system. DNA breaks at tandem I-SceI sites release the puromycin resistance gene, allowing NHEJ repair to join a promoter to GFP expressing sequence. **(E)** After generation of a stable U2OS clone containing the NHEJ reporter construct, *RB1* was deleted as above and confirmed by western blotting. **(F)** NHEJ repair efficiency was determined by transfecting an I-SceI endonuclease expression vector (I-SceI +), or negative control, and PI- and GFP+ cells were quantitated by flow cytometry (n=3). **(G)** Cells were treated with 2 Gy IR and fixed 1 hr or 24 hr after treatment and stained for γH2AX. Untreated cells were stained in parallel. Three clones per genotype were used and γH2AX foci were quantified. The proportion of cells with discrete numbers of foci are shown as histograms, while the cumulative frequency of foci are shown as inset. Differences in foci distribution were determined using the Kolmogorov-Smirnov test. A green asterisk indicates *RB1*^−/−^ is statistically different than the other genotypes, while a black asterisk indicates all genotypes are statistically significantly different from each other. All error bars are 1 SEM. \**P* < 0.05.

To assess how *RB1* mutant U2OS cells respond to DNA breaks, each genotype was exposed to 2 Gy of gamma radiation (γIR). Cells were fixed and stained for γH2AX and DAPI at various time points to measure the amount of DNA damage. One hour after γIR, there was a pronounced increase in γH2AX foci in all genotypes compared to untreated cells (Fig. 3G). However, the amount of DNA damage was significantly greater in *RB1*^−/−^ clones compared to heterozygous or wild type cells. After 24 hr, most of the DNA damage was repaired, and the *RB1*^−/−^ still had more γH2AX foci than the other two genotypes. Overall, this indicates that cells completely lacking RB are more sensitive to γIR, likely because they are not able to repair DNA breaks as efficiently by HR repair.

Next, we investigated the fidelity of mitosis to determine if the elevated levels of DNA damage and impaired HR repair impacted chromosome segregation and aneuploidy (Gelot et al., 2015). Flow cytometry was performed on *RB1* deficient cells that were labeled and stained with BrdU and propidium iodide (Cecchini et al., 2012). This analysis failed to show statistically different changes in cell cycle phases between the different genotypes (Fig. 4A). However, when DNA content greater than 4N was analyzed, there was a significant difference between wild type and *RB1* knockout cells, with *RB1*^+/−^ cells showing an intermediate value (Fig. 4B). This suggests mitotic errors in these cells may lead to aneuploidy.

**Figure 4.**
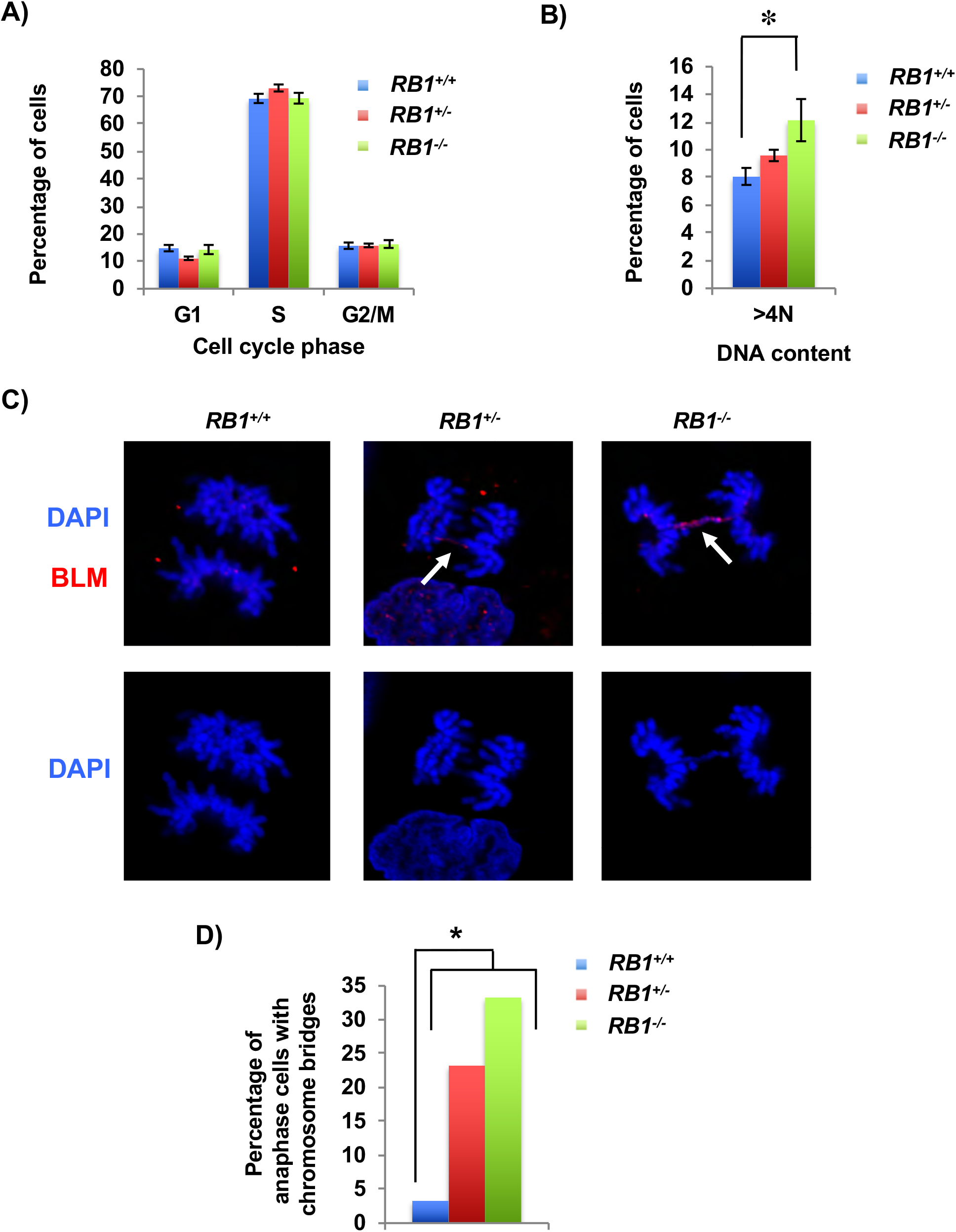
Increased mitotic errors in *RB1* mutant cells. **(A)** BrdU and propidium iodide staining followed by flow cytometry were used to determine cell cycle phase distribution of asynchronous cultures. Four clones for the wild type and knockout genotypes and three clones for the heterozygous genotype were analyzed. **(B)** Flow cytometry analysis shows the proportion of cells with greater than 4N DNA content. Means were compared using a one-way ANOVA). **(C)** Cells in anaphase were imaged by fluorescence microscopy using DAPI (blue) and BLM (red) in cells from each *RB1* genotype. Arrows indicate anaphase bridges that are stained by both DAPI and BLM. **(D)** The number of anaphase cells with DAPI stained chromosome bridges were quantitated. The proportion of cells with bridges is significantly higher in the knockout and heterozygous clone compared to the wild type clone as determined by the χ^2^-test. All error bars are ±1 SEM. \**P* < 0.05.

To further investigate mitotic defects, cells were stained with DAPI and antibodies to BLM to visualize chromosome bridges, and mitotic figures were imaged using confocal microscopy (Fig 4C). We observed abundant chromosome bridges in *RB1*^−/−^ and *RB1*^+/−^ mutant cells (Fig. 4D). In the *RB1* mutants there were some ultra-fine bridges (UFBs), which are “thread-like” DNA structures that stain only with BLM (Chan and Hickson, 2011). However, the majority of BLM bridges stained with DAPI, indicating that anaphase bridges were most common (SFig. 5). Anaphase bridges are known to occur in homologous recombination (HR)-defective cells, while UFBs can be induced by replication stress (Gelot et al., 2015). Our analysis of mitotic errors is consistent with a defect in HR repair being the source of chromosome bridges in anaphase of these *RB1* mutant cells.

### Increased lung metastases in *RB1* mutant xenografts

To further characterize the effects of induced *RB1* mutations in cells that already possess RB-pathway defects, we performed xenograft experiments to determine if new cancer relevant properties arise upon loss of RB. We injected cells subcutaneously into immune compromised mice and allowed tumors to form for an eight-week period before analyzing growth by mass and histology (Fig. 5A). This analysis revealed a highly cellular structure with abundant mitotic figures. Cells appeared epithelioid with no definite features of osteoid differentiation and they had small areas of glandular organization, this phenotype was consistent among all genotypes (Fig. 5B). Tumor masses were determined at end point and were not statistically different between genotypes, with *RB1*^−/−^ even trending towards a smaller size (Fig. 5C). Mice were also tail-vein injected and cell dissemination and proliferation were allowed to proceed for eight weeks at which time lungs were harvested, fixed, and sectioned. Hematoxylin and eosin-stained sections were digitally analyzed to quantitate cellular infiltration of U2OS cells (Fig. 5D). This revealed a striking increase in *RB1*^−/−^ U2OS cells in the lungs of these mice compared to control and *RB1*^+/−^ cells (Fig. 5E). There were significantly more individual clusters of *RB1*^−/−^ cells per lung than the other genotypes (Fig. 5F), further suggesting that *RB1* loss increased the efficiency of dissemination or establishment in the lung. Lastly, the percentage of tumor cell area per cluster of cells is lower in both *RB1* mutant genotypes indicating that control cells form rarer, larger clusters of cells, whereas *RB1* mutants tend to seed more efficiently and perhaps proliferate more slowly (Fig. 5G). We note that one mouse injected with *RB1*^+/−^ cells showed extensive dissemination that was highly reminiscent of *RB1*^−/−^ (SFig. 6A&B). PCR analysis of tumor material from this mouse confirmed that these cells maintained their *RB1*^+/−^ genotype, as the remaining wild type *RB1* allele was not lost (SFig. 6C).

**Figure 5.**
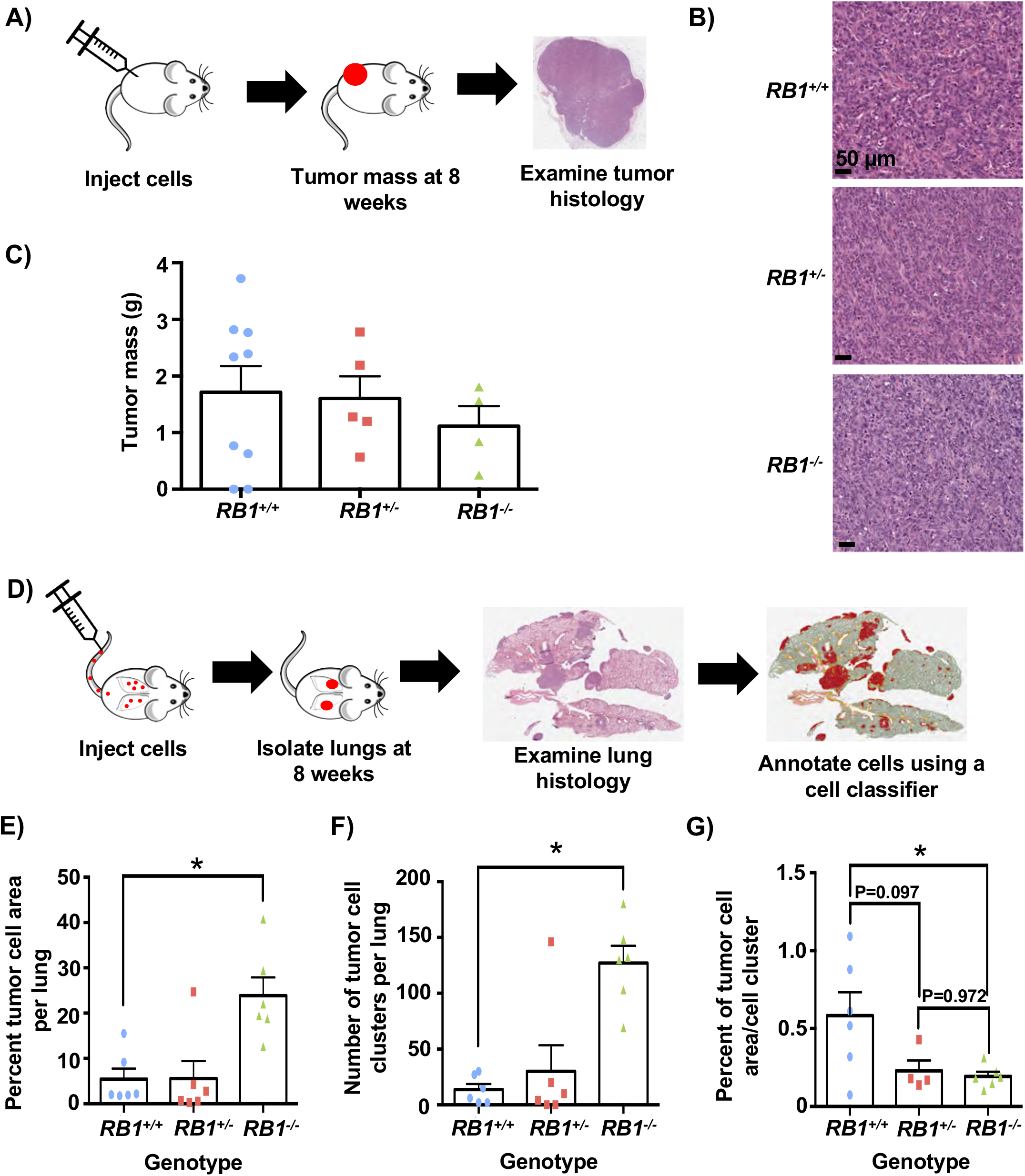
Mouse xenografts reveal increased lung metastases with *RB1* mutant cells. **(A)** Illustration of analysis of subcutaneous injections of *RB1* mutant U2OS cells. Tumors were allowed to form for just under eight weeks before analyzing tumor mass and histology. **(B)** Representative H&E stained tissue sections from each genotype of tumor. **(C)** Tumor masses from subcutaneous injected cells are shown. **(D)** Schematic of tail vein injections to study dissemination to lung. Mice were injected and cell dissemination and proliferation were allowed to proceed for eight weeks. Lungs were then isolated, sectioned and stained with H&E, and analyzed using QuPath. **(E)** The percentage of tumor cell area was calculated from lung sections. **(F)** Tumor cell clusters were counted using the assistance of this digital classifier. **(G)** Percent tumor cell area was divided by the number of clusters to determine the average tumor cell area per cluster. All error bars are +1 SEM. \**P* < 0.05.

These xenograft experiments indicate that *RB1* mutations in U2OS cells have little effect on growth rate of primary tumors. Interestingly, *RB1*^−/−^ cells are much more efficient in colonizing recipient mouse lungs, suggesting that in addition to the DNA damage and genome instability phenotypes described earlier, *RB1*loss in RB-pathway deficient cells imparts characteristics that enable cancer progression.

## Discussion

To test if *RB1* loss is important in cancer cells that already possess mutations disrupting the RB-pathway, we used CRISPR/Cas9 to create non-functional *RB1* alleles. Using γH2AX staining of cells, we found that even within cells deficient for one copy of *RB1*, there is an increase in spontaneous DNA damage. Experiments that probe drug sensitivity of *RB1* mutant cells, along with ChIP-Seq analysis of γH2AX foci and analysis of DNA damage repair pathways, suggest a complex picture of cellular defects. We did not detect specific hotspots of DNA damage, but sensitivity to peroxide and cisplatin suggest oxidative damage may cause sporadic DNA damage as reactive oxygen species are elevated in *RB1* deficient cells. Mutant *RB1* cells also have more abnormal mitoses characterized by anaphase bridges, and reporter assays detect defects in HR repair suggesting another endogenous source of damage. Overall, this study reveals that *RB1* mutations lead to increased DNA damage that may enhance cancer progression.

The DNA damage phenotype caused by either single copy or homozygous mutation to *RB1* is unlikely to be attributable to a single root cause and we expect that *RB1* deficiency may affect DNA damage in different cancer cells in varied ways. We report that *RB1* deletion compromises HR-repair but not NHEJ and this is consistent with one previous report (Velez-Cruz et al., 2016), but contradicts another (Cook et al., 2015). Given that we observe more than 80% of U2OS cells incorporating BrdU in a brief pulse, they are likely biased towards HR repair pathways. This is consistent with γH2AX foci not being accompanied by 53BP1, and the activity levels of NHEJ reporters being almost an order of magnitude less than HR values, suggesting that U2OS cells are primed to use the HR repair pathway and thus phenotypes in these cells reflect this reality. We expect that DNA damage and defective repair described in this report are relevant to cancer progression phenotypes because graded differences between *RB1* wild type, heterozygous, and homozygous genotypes are reflected in 8-oxoguanine abundance, aneuploidy, and anaphase bridges. This stepwise trend in severity of phenotype is similarly evident in the behaviour of *RB1^+/+^, RB1^+/−^*, and *RB1*^−/−^ mutant tail vein xenograft experiments.

Single copy loss of *RB1* may contribute to cancer in a number of ways. Primary *RB1*^+/−^cells from a number of sources are prone to mitotic errors (Coschi et al., 2014; Gonzalez-Vasconcellos et al., 2013; Zheng et al., 2002), and precursor lesions to Retinoblastoma are characterized by aneuploidy (Dimaras et al., 2008), suggesting this is an early step in this disease. Therefore, partially defective *RB1* can contribute to the early stages of cancer through a distinct set of effects. However, a number of studies have highlighted that *RB1* loss is statistically enriched in advanced stages of cancer progression (Beltran et al., 2016; McNair et al., 2017; Robinson et al., 2017). In addition, analysis of the landscape of cancer alterations in TCGA consortia reveals so called ‘shallow deletions’ of *RB1* as relatively commonplace (Fig. 6), and these events are unlikely to be explained by random, unselected events (Beroukhim et al., 2010). From this perspective, our study offers critical proof of concept that late stage loss of one copy of *RB1* can create cancer enabling phenotypes in the host cell, even if it already possesses RB-pathway mutations. Complete elimination of *RB1* has a stronger effect on DNA damage phenotypes and dissemination to the lungs, as demonstrated in this study, however, highly abundant single copy *RB1* loss may represent a mutational compromise in which advantageous phenotypes are acquired with an economy of mutational changes (Davoli et al., 2013).

**Figure 6.**
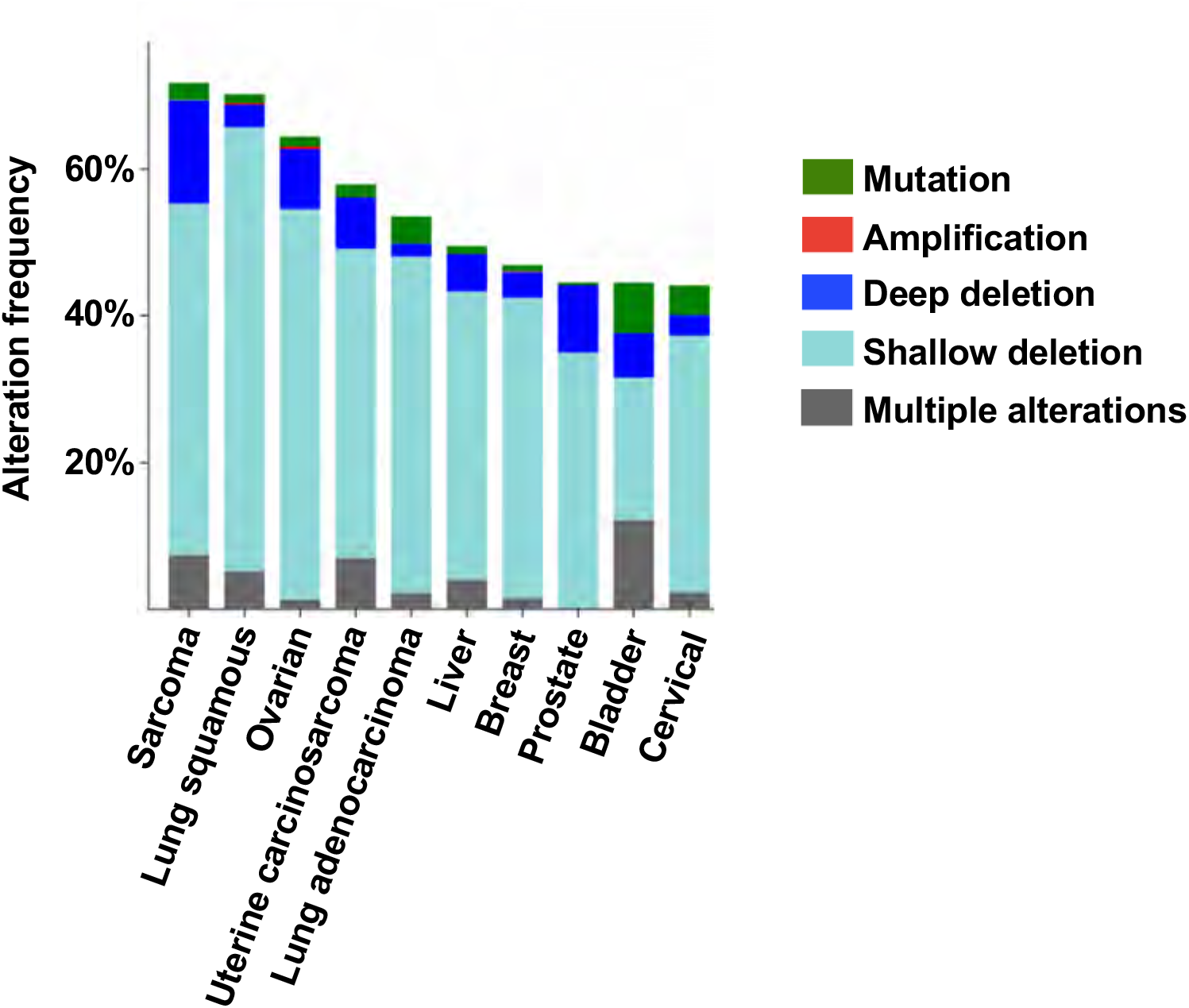
The most common *RB1* alteration in cancer are shallow deletions. The 10 most prevalent cancers were analyzed for *RB1* gene alterations using TCGA and Pan-Cancer Atlas data using cBioPortal. A deep deletion is consistent with biallelic loss of *RB1*, whereas a shallow deletion is suggestive of heterozygous *RB1* deletion, but could be due to stroma contributions or cancer heterogeneity.

A number of studies correlate absence of RB expression with improved patient survival following treatment that includes platinum based chemotherapy (Cecchini et al., 2015; Garsed et al., 2017). Loss of RB correlated with improved survival of lung adenocarcinomas treated by resection and adjuvant cisplatin or carboplatin, and a vinca alkaloid (Cecchini et al., 2015). More recently patients with high-grade serous ovarian cancer (HGSC) that experienced exceptionally good clinical outcomes were studied (Garsed et al., 2017). These patients were treated with platinum-based agents and loss of RB was associated with long-term survival. From this perspective, U2OS cells engineered to be deficient for *RB1* demonstrate that RB loss increases sensitivity to cisplatin. Given that *RB1* mutant cells were not more sensitive to another agent that induces DNA breaks, etoposide, we interpret *RB1* deficiency to create a unique sensitivity to cisplatin that may relate to defective HR and higher endogenous ROS levels that create a highly specific sensitivity to this class of chemotherapy.

In conclusion, although there are many cancers that have mutations in the RB-pathway that spare the *RB1* gene itself, further mutations to *RB1* surprisingly create cancer relevant characteristics that may influence disease progression. The frequency of shallow deletions of *RB1* across many cancers suggests that disease progression may select for these characteristics.

## Acknowledgements

The authors wish to thank colleagues for discussions and encouragement over the course of this study. AEM was a recipient of fellowship support from NSERC and OGS. This work was supported by funding from the Cancer Research Society, and the CIHR (MOP 324579). FAD is the Wolfe Senior Fellow in Tumor Suppressor Genes at Western University.

## Author Contributions

MVR, DTP, and MCD carried out experiments. CVH interpreted experimental data. ACC and JS contributed reagents and assisted in writing the manuscript. AEM and FAD conceived of the project, carried out experiments, and wrote the manuscript.

## Methods

### Cell Culture

U2OS cells and the resulting clones were grown in Dulbecco’s modified Eagle’s medium (DMEM) supplemented with 10% fetal bovine serum (FBS), 2 mM L-glutamine, 50 U/mL penicillin and 50 μg/mL streptomycin. H460 and H1792 cells and the resulting clones were grown in Roswell Park Memorial Institute (RPMI) 1640 medium supplemented with 10% FBS, 2 mM L-glutamine, 50 U/mL penicillin and 50 μg/mL streptomycin. Cells were grown at 37°C in humidified air containing 5% CO2.

### Generation of *RB1* deletions using CRISPR

For creation of *RB1* deletions, single guide RNAs (sgRNAs) targeting exon 22 of *RB1* were selected by using the CRISPR Design tool at http://crispr.mit.edu/(Cong et al., 2013). The sgRNA sequences were as follows: 5’-CACCGTATTATAGTATTCTATAACT-3’ (X22B-top), 5’-AAACAGTTATAGAATACTATAATAC-3’ (X22B-bottom), 5’- CACCGAGGATACTTTTGACCTACCC-3’ (X22C-top), 5’- AAACGGGTAGGTCAAAAGTATCCTC (X22C-bottom). The X22B and X22C guides were each cloned into the pX459 plasmid (with wild type Cas9; Addgene #48139) and the pX462 plasmid (with the D10A mutant version of Cas9; Addgene #48141) Both plasmids contain a puromycin resistance cassette. Cells were seeded at a density of 10^5^ cells per well in a 6-well dish, and the next day a total of 1 μg per well of a 1:1 mix of X22B and X22C CRISPR plasmids (either pX459 or pX462) was transfected by use of X-tremeGENE HP DNA transfection reagent (Roche). The next day, each well was replated onto a 15 cm dish, and the day after that cells were cultured in selection medium with 2 μg/mL of puromycin for 2 days. Following that, cells were grown in normal cell culture media for approximately 12 days, following which single colonies were picked from the 15 cm plates using mechanical detachment with a pipette tip and placed into wells of a 48-well dish and allowed to grow. Cells were further passaged onto larger plates, and were genotyped using the following primers: X22F primer, 5’-TTACTGTTCTTCCTCAGACATTCAA-3’; and X22R primer, 5’- GGATCAAAATAATCCCCCTCTCAT-3’. PCR products (445 bp for the wild type band) were run in an agarose gel, individual bands were gel purified using a Sigma GenElute Gel Extraction kit and sent for Sanger sequencing using the X22F and X22R primers shown above. For clones with multiple PCR products, bands were purified and sent for sequencing separately to determine individual *RB1* alleles in each clone. For alleles that could not easily be resolved by gel electrophoresis, PCR products were cloned into vectors using either the TOPO TA Cloning Kit for Sequencing (Invitrogen) or the CloneJET PCR Cloning Kit (Thermo Scientific).

The top scoring off-target intragenic locations determined for each gRNA using the CRISPR Design tool were also sequenced to probe for mutations. gRNA X22B had a potential off-target site in *ZNF699* (X22B_OT_ZNF699_F, 5’-GTGCCCTAAAACACTGAGGGA-3’; and X22B_OT_ZNF699_R, 5’-TTTATGATCAACAAGGACCAGAGC-3’) while X22C has a potential off-target site *ALDH1L1* (X22C_OT_ALDH1L1_F, 5’- GCCACGCTATGCTTGTGATG-3’; and X22C_OT_ALDH1L1_R, 5’- CACCCCAGAGAAGGGAACAC -3’). PCR products were gel purified as above and sent for Sanger sequencing using their respective primers.

Nuclear extracts were prepared from U2OS CRISPR clones of interest and western blotting was carried out using previously described protocols (Cecchini and Dick, 2011). Antibodies raised against RB (clone G3-245, BD Pharmingen; C-15, Santa Cruz) and Sp1 (H-225, Santa Cruz) were used for western blotting. Samples were western blotted using standard techniques.

To generate additional *RB1* knockout and control cell lines, sgRNAs targeting either exon 2 of *RB1* (5’-GGAGAAAGTTTCATCTG-3’) or a gene desert region of the genome (5’- TGAGCCTATATTAATTGG-3’) were utilized. The sgRNAs were cloned into the lentiCRISPR v2 vector (Addgene #52961), which also encodes Cas9. To generate lentivirus, 293T cells were transfected with the sgRNA vector and a 1:1:1 mixture of lentiviral packaging constructs (Addgene #12251, #12253, #8454) using polyethylenimine transfection reagent. Twenty-four hours after transfection, the 293T media was replaced, and recipient cells (U2OS, NCI-H1792, NCI-H460) were seeded for infection. The following day, media on the recipient cells was replaced with lentiviral media, and polybrene was added at a final concentration of 8 μg/mL. A second infection was performed the next day. Infected cells were then selected with 2 μg/mL puromycin for 3 days. To generate isogenic clones, populations of knockout (or control) cells were FACS sorted as single cells in 96-well plates (BD FACSAria II) and allowed to grow for approximately 2 weeks. Colonies were then expanded and screened for loss of RB by immunoassay using the Simple Western™ system according to the manufacturer’s instructions. Successful knockout clones were also genotyped to confirm clonogenic origin. Genomic DNA was extracted using the PureLink Genomic DNA Mini Kit (Invitrogen), and the region surrounding the cut site was amplified by PCR using the following primers: X2F, 5’- TCACAGAAGTGTTTTGCTGCTT-3’; X2R, 5’-TTTGGTGGGAGGCATTTATGGA-3’. PCR products were purified using the DNA Clean and Concentrator Kit (Zymo Research) and sent for sequencing.

### Fluorescence Microscopy

Cells grown either on glass coverslips or in glass bottom plates were fixed in phosphate-buffered saline (PBS) containing 4% paraformaldehyde for 10 min and then permeabilized with PBS-0.3% Triton X-100 for 10 min at room temperature. The fixed cells were blocked in blocking buffer (PBS-0.3% Triton X-100 with either 5% donkey or goat serum depending on the species in which the secondary antibodies were raised) for at least 1 h at room temperature. Cells were then incubated with primary antibody in blocking buffer at room temperature for 1 h or at 4°C overnight. Antibodies raised against RB (clone G3-245, BD Pharmingen), γH2AX (clone JBW301, EMD Millipore), 53BP1 (H-300, Santa Cruz) and BLM (C-18, Santa Cruz) were used for IF. After 3 washes with PBS-0.3% Triton X-100, cells were incubated with secondary antibody diluted in blocking buffer for 1 h at room temperature. Cells were washed twice with PBS-0.3% Triton X-100, incubated with 100 ng/mL 4’,6-diamidino-2-phenylindole (DAPI) in PBS-0.3% Triton X-100 for 5 min, washed twice more with PBS-0.3% Triton X-100 and then washed once with PBS before mounting with Slowfade Gold Antifade mountant (S36936, ThermoFisher Scientific).

For 8-oxoG staining, cells were fixed and blocked as above, then washed with PBS-0.3% Triton X-100 and incubated in RNase solution (0.2 mg/mL RNase A, 10 mM Tris-HCl (pH 7.5), 15 mM NaCl, 0.1% Triton X-100 in 1X PBS) for 1 h at room temperature. Cells were washed with PBS-0.3% Triton X-100 and then incubated in 2 M HCl for 10 min at room temperature, followed by a rinse with 50 mM Tris-HCl (pH 8.0). Cells were washed with PBS-0.3% Triton X-100, and then primary antibody incubation, using α-DNA/RNA Damage antibody raised against 8-oxoG (clone 15A3, ab62623, Abcam) and all subsequent steps were completed as above.

For confocal microscopy of RB, γH2AX and BLM, cells were examined on an Olympus Fluoview FV1000 confocal microscope system. For confocal microscopy of 53BP1, a Nikon A1R confocal microscope was used. For non-confocal microscopy, images were acquired using a Zeiss Axioskop 40 microscope and Spot flex camera. Foci were quantified using the Focinator (Oeck et al., 2015), while overall staining intensity in cells was quantified by ImageJ (Schneider et al., 2012).

### Gamma irradiation of cells

Cells subjected to γIR were plated at 100,000 cells per well in 6 well dishes with glass coverslips on the bottom. The next day, cells were exposed to a cobalt 60 source until a dose of 2 Gy was received. Cells were placed back in the cell culture incubator until the appropriate time point after treatment to fix cells for IF.

### NHEJ and HR repair assays

For the HR repair assay, pDRGFP was used which was a gift from Maria Jasin (Addgene plasmid #26475; http://n2t.net/addgene:26475; RRID:Addgene_26475) and for the NHEJ assay, pimEJ5GFP was used which was a gift from Jeremy Stark (Addgene plasmid #44026; http://n2t.net/addgene:44026;RRID:Addgene_44026). pDRGFP was linearized using EcoRV and pimEJ5GFP was linearized using XhoI. These linearized fragments were then individually used for transfection using Lipofectamine 3000 transfection reagent (Invitrogen) into U2OS cells. The next day, each well was replated onto a 10 cm dish, and a day later, cells were cultured in selection medium with 2 μg/mL of puromycin for 3 days. To isolate single cell colonies, limiting dilutions were then used to seed cells into 96 well plates. After approximately 3 weeks, wells with growth from single cell isolates were transferred to single wells of 12 well plates and after a few days were treated with puromycin again to ensure the selected clones did still contain either the NHEJ or HR constructs.

To determine the reporter efficiency in the isolated clones, 2 sets of transfections were performed per clone, again using Lipofectamine 3000 transfection reagent (Invitrogen). For the first set of transfections, each clone was transfected with a plasmid expressing the I-SceI endonuclease, pCBASceI, which was a gift from Maria Jasin (Addgene plasmid #26477; http://n2t.net/addgene:26477; RRID:Addgene 26477), and a blasticidin marker, pMSCV-Blasticidin, which was a gift from David Mu (Addgene plasmid #75085; http://n2t.net/addgene:75085; RRID:Addgene_75085). The second set of transfections was with an empty backbone plasmid, pCAG-FALSE, which was a gift from Wilson Wong (Addgene plasmid #89689; http://n2t.net/addgene:89689; RRID:Addgene_89689) and pMSCV-Blasticidin. For both of these transfections, the blasticidin resistance plasmid was used in a 1:3 ratio with the complementary plasmid. The next day, each well was replated onto a 10 cm dish, and the day after that cells were cultured in selection medium with 10 μg/mL of blasticidin for 1 week. GFP positive cells were then quantified by flow cytometric analysis (FACS). To prepare cells for FACS, they were washed with PBS, trypsinized, resuspended in culture media, and washed twice with PBS. Cell pellets were then resuspended in 0.5 mL of flow cytometry staining buffer with propidium iodide (0.05% sodium azide and 0.5% BSA in 1X PBS with 0.01 mg/mL propidium iodide). For each reporter construct, the clone with the highest ratio of GFP signal when transfected with pCBASceI to GFP signal when transfected with pCAG-FALSE was selected for future studies.

To introduce CRISPR constructs into selected clones for each repair reporter, lentivirus particles were generated in HEK293T cells. Lentivirus was created for both lentiCRISPR v2 with no guide RNA inserted, and for lentiCRISPR v2 with the X22B sgRNA sequences for *RB1* (from above) inserted. lentiCRISPR v2 was a gift from Feng Zhang (Addgene plasmid #52961; http://n2t.net/addgene:52961; RRID:Addgene_52961). The X22B *RB1* guide sequences were inserted into the lentiCRISPR v2 plasmid as previously described (Sanjana et al., 2014; Shalem et al., 2014). Culture media containing lentiviral particles were transferred to appropriate U2OS HR and NHEJ reporter clones for 48 hours, followed by selection with 4 μg/mL puromycin for at least 5 days.

U2OS HR and NHEJ reporter clones that were infected with lentiCRISPR v2 plasmids, either with or without the *RB1* guide, were then transfected with both pCBASceI and pMSCV-Blasticidin or pCAG-FALSE and pMSCV-Blasticidin and selected, as above, and analyzed by FACS to determine repair efficiency. Cells grown in parallel to the transfected cells were used to prepare nuclear extracts for western blotting.

### Determination of IC50 concentrations

For IC50 assays, cells were seeded at a density of 12,000 cells per well in 96 well dishes. Twenty-four hours after plating cells, media was replaced with media containing the drugs of interest at the appropriate concentrations. Technical triplicates were analyzed for each biological replicate. Serial dilutions of stock solutions of aphidicolin (APH), hydrogen peroxide (H_2_O_2_), etoposide, hydroxyurea (HU) and cisplatin were created so that a constant amount of drug was added to the media for each drug concentration used. After 72 h, alamarBlue was added to an amount equal to 10% of the volume in the well (i.e. 10 μL per well with 100 μL of media and drug). After 4 h of incubation, cytotoxicity was measured using a Synergy H4 Hybrid Reader (BioTek, USA) using excitation/emission wavelengths of 560 nm/590 nm. Values were corrected using a blank of media and alamarBlue only. The amount of fluorescence of alamarBlue for each well of drug treated cells was then normalized to the fluorescence value obtained for the untreated cells of the same technical replicate. These normalized fluorescence values relative to untreated cells were then analyzed using Prism. The drug concentrations were log transformed and the data was subsequently fit to a curve using nonlinear regression (log(inhibitor) vs. response (three parameters)). IC50 values were obtained from the best fit values, and IC50 values from three biological replicates were compared using Ordinary one-way ANOVA and Tukey’s multiple comparisons test.

### ChIP-sequence

ChIP was conducted according to protocols adapted from Cecchini *et al.* (Cecchini et al., 2014). Briefly, cross-linked chromatin was sonicated so most chromatin was ≤400 bp. Sheared chromatin was then normalized between experimental groups and pre-cleared with protein G Dynabeads and IgG. Pre-cleared chromatin was then incubated with protein G Dynabeads and ChIP antibodies to immunoprecipitate proteins. Antibodies raised against γH2AX (clone JBW301, EMD Millipore) and H4 (clone 62-141-13, EMD Millipore) were used for ChIP. Cross-links were reversed at 65°C, and samples were treated with RNase and proteinase K. DNA was isolated for library preparation, and 20 replicates per genotype for γH2AX ChIP-Seq were pooled to achieve DNA yield required for library preparation (NEBNext Ultra II DNA Library Prep Kit). ChIP libraries were sequenced using an Illumina NextSeq (High output 75 cycle kit), and processed reads are deposited in GEO (GSE125379).

Resulting FASTQ reads were aligned to the human genome build hg19 using Bowtie2 version 2.3.0 (Langmead and Salzberg, 2012). The following command was used: bowtie2 -t -p 4 -D 15 -R 2 -L 32 -i S,1,0.75 -x hg19 -U <reads>.fastq -S <output>.sam. Peaks were identified using MACS2 version macs2 2.1.1.20160309 according to parameters stated below and the options to detect broad peak distributions for histone marks (Zhang et al., 2008). For H4 ChIP-Seq, the corresponding inputs were used as the control and for γH2AX ChIP-Seq, the first input replicate was used as the control. The following command was used: macs2 callpeak -t <ChIP>.bam -c <input>.bam -n <output> --outdir ./macs2/ -g hs --broad --broad-cutoff 0.1.

To find abundance of ChIP-Seq reads in common fragile sites (CFS), the cytogenetically determined locations of CFS, as determined previously by Lukusa and Fryns (2008), were converted to human genomic coordinates (hg19) using the UCSC Genome Browser (Lukusa and Fryns, 2008; Tyner et al., 2017). Bedtools coverage was then used to find the number of alignments for each ChIP-Seq sample within the individual CFS (Quinlan, 2014). The abundance of reads that mapped to CFS were then converted to proportions by dividing by the total number of mapped reads. The proportion data were further normalized against input control and then ratios were made comparing the mutant proportions to the wild type proportions. A two-tailed one-sample t-test was performed to test if the normalized mean read count proportions of the *RB1*^+/−^ and the *RB1*^−/−^ ChIP-Seq assays at each of the CFS is equal to the normalized read count proportion of the corresponding CFS from the wild type control. A multi-test correction was applied to the calculated P-values (using “fdr” method from “p.adjust” function in R). Statistical analysis of sequence data was performed using R (version 3.4.2) and the plotting function used was lattice (v0.20-35).

For repeat analysis, another set of alignments were performed. For this analysis, reads were mapped using Bowtie version 1.2.1.1 with high stringency to the hg19 genome (Langmead et al., 2009). The following command was used: bowtie -S --best -m 1 --chunkmbs 500 -p 4 -t -un <not_aligned_unique> --max <multiple_reads_unique> hg19 <reads>.fastq <output>.sam.

All remaining reads were mapped to repeat containing indexes using previously reported methods (Day et al., 2010), with indices also being derived from Repbase and Tandem Repeats Databases (Bao et al., 2015; Gelfand et al., 2007). For these remaining repeat alignments, the –m 1 parameter of the Bowtie mapping was changed to –k 1. Finally, all remaining reads were remapped to hg19 at low stringency to exhaustively match sequence tags to the mouse genome.

The abundance of sequence tags that mapped to non-unique regions of the genome were compared by using log2 ratios of γH2AX precipitable tags per million mapped reads in mutant versus wild type and converted into heat maps using matrix2png (Pavlidis and Noble, 2003). To test for significance of enrichment of reads mapped to various repeat categories, the same analysis to test for significance within CFS was used (see above).

### Flow Cytometry

Cells were plated on 6 cm plates at a density of 100,000 cells per plate. Approximately 24 h after, cells were pulsed with BrdU for a duration of 30 min. Cell cycle analysis was then carried out as previously described (Cecchini et al., 2012).

### Measurement of Reactive Oxygen Species

Cells were plated in 96 well plates at a density of 12,000 cells per well in DMEM without phenol red (31053-028, ThermoFisher Scientific). H_2_O_2_ or cisplatin were added 24 h after seeding cells. Seventy-two hours later, 5(6)-carboxy-2’,7’-dichlorodihydrofluorescein diacetate (carboxy-H2DCFDA; CA-DCF-DA; (C400, ThermoFisher Scientific)) at a stock concentration of 20 mM in DMSO was diluted in DMEM without phenol red to a concentration of 40 μM. This master mix of media and CA-DCF-DA was added directly to the wells already containing media and the drug of interest, to obtain a final concentration of 20 μM CA-DCF-DA. The cells were then put in the incubator for 45 min and readings were obtained using a Synergy H4 Hybrid Reader (BioTek, USA) using excitation/emission wavelengths of 492 nm/525 nm. Technical triplicates were analyzed for each biological replicate and the average background readings (cells treated with the highest concentration of the drug of interest for 72 h and DMSO in place of CA-DCF-DA) from each cell line were subtracted from the average of each treatment reading for analysis of fluorescence.

### Mouse xenografts

U2OS clones were grown in cell culture to approximately 80% confluence. Cells were washed with PBS, trypsinized, centrifuged and washed 3 times with Hanks’ Balanced Salt Solution (HBSS, 1X). Cells were then resuspended in HBSS at a concentration of 5×10^6^ cells/mL so that 200 μL contained the 1×10^6^ cells required for each injection.

For subcutaneous injections, mice were approximately 8 weeks old and for the tail vein injections mice were approximately 13 weeks old when injected. All mice were given at least 3 days to acclimatize. All mice were female NOD.Cg-*Prkdc^scid^Il2rg^tmlWjl^*/SzJ (stock number 005557, The Jackson Laboratory) and were housed and handled as approved by the Canadian Council on Animal Care, under an approved protocol (2016-068).

For subcutaneous injections, 1×10^6^ *RB1*^+/+^ cells were injected into the left flank of all mice, and 1×10^6^ *RB1*^+/−^ or *RB1*^−/−^ cells were injected into the right flank. The mice used for the subcutaneous injections were euthanized approximately 8 weeks after injection. Necropsies were performed and tumor mass was determined. Tumors were then fixed in formalin for 48 hr and processed for histological assessment.

For tail vein injections, 1 × 10^6^ cells were injected into the lateral tail vein. Mice were euthanized 8 weeks after tail vein injection. All animals were subjected to a thorough necropsy and all lungs as well as any abnormal tissues or organs were fixed in formalin for 48 hr and processed for histological assessment.

All tissues of interest from both studies were embedded, sectioned and stained with hematoxylin and eosin according to standard methods. Slides were imaged using an Aperio ScanScope slide scanner (Leica Biosystems).

For quantitative pathology of lungs from tail vein injected mice, images were analyzed using QuPath (Bankhead et al., 2017). Briefly, annotations were drawn around each individual lung. Within these annotated lungs, cells were detected using the cell detection command. The features within these cells were then smoothed by using the add smoothed features command (using 25 μm as the radius). Within the lungs, regions containing different cell types were annotated and these annotations were used to train a cell classifier. All possible 67 cell features were used to the build the random trees classifier, using default parameters. A script was then made to determine the total cell area of all cell types called by the classifier within each lung, and the percentages of tumor cell area was calculated from these values. Tumor cell clusters were manually counted using the cell types determined by the classifier; anything thought to have derived from a single cell seeding event was considered a tumor cell cluster.

To determine the *RB1* genotype of seeded U2OS cells of interest, embedded mouse lung tissue was deparaffinized, lysed, formalin crosslinks were reversed, and DNA was isolated according to manufacturer’s instructions (QIAamp DNA FFPE Tissue Kit, Qiagen). DNA was genotyped as above using PCR using genotyping primers (X22F and X22R).

### Data extraction from cBioPortal

Only TCGA studies used for the Pan-Cancer Atlas with 150 samples or more on cBioPortal were selected to query (Cerami et al., 2012; Gao et al., 2013). Data from cBioPortal was obtained in January 2019. Mutation and CNA data was analyzed, with the gene set user-defined list being entered as “RB1: AMP HOMDEL HETLOSS mut”.

## Supplemental Figures

**Supplemental Figure 1.**
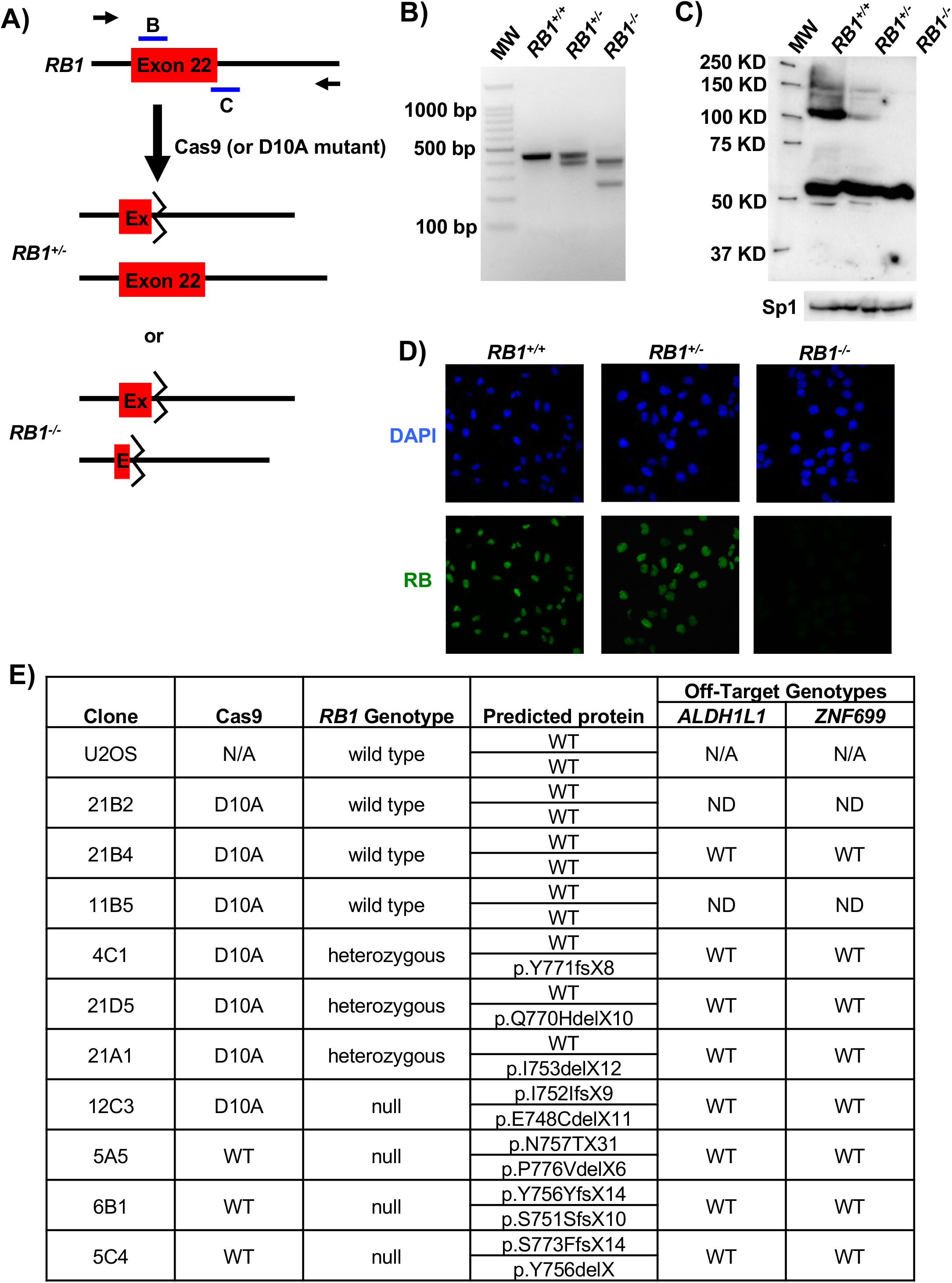
Strategy to generate *RB1* mutant U2OS cells. **(A)** Schematic depicting the relative locations of gRNA pairs (denoted as B and C) used in this study in relation to exon 22 coding sequence. The locations of flanking primers used to detect deletions are represented as arrows. Hypothetical heterozygous and homozygous *RB1* mutant gene structures containing deletions of exon 22 that are expected following transfection of gRNA and Cas9 encoding plasmids are shown. **(B)** An ethidium bromide stained, agarose gel shows examples of wild type, heterozygous, and homozygous mutant *RB1* genotypes that are detected by PCR amplification of exon 22 sequences. **(C)** A representative western blot showing pRB expression in control, heterozygous, and homozygous mutant cells is shown on top. SP1 loading control is shown on the bottom. **(D)** Immunofluorescence microscopy was used to detect RB expression in cultures of control, heterozygous, or homozygous mutants (shown in green). Cells were counterstained with DAPI to visualize nuclei (blue). **(E)** Table summarizing the characterization of the U2OS *RB1* mutant clones used in this study. Genotypes were determined by PCR and sequencing, and amino acid coding changes were predicted based on nucleotide sequences. The top scoring off-target intragenic locations determined for each gRNA, *ZNF699* for X22B and *ALDH1L1* for X22C, were also sequenced to probe for unwanted mutations. ND=not determined.

**Supplemental Figure 2.**
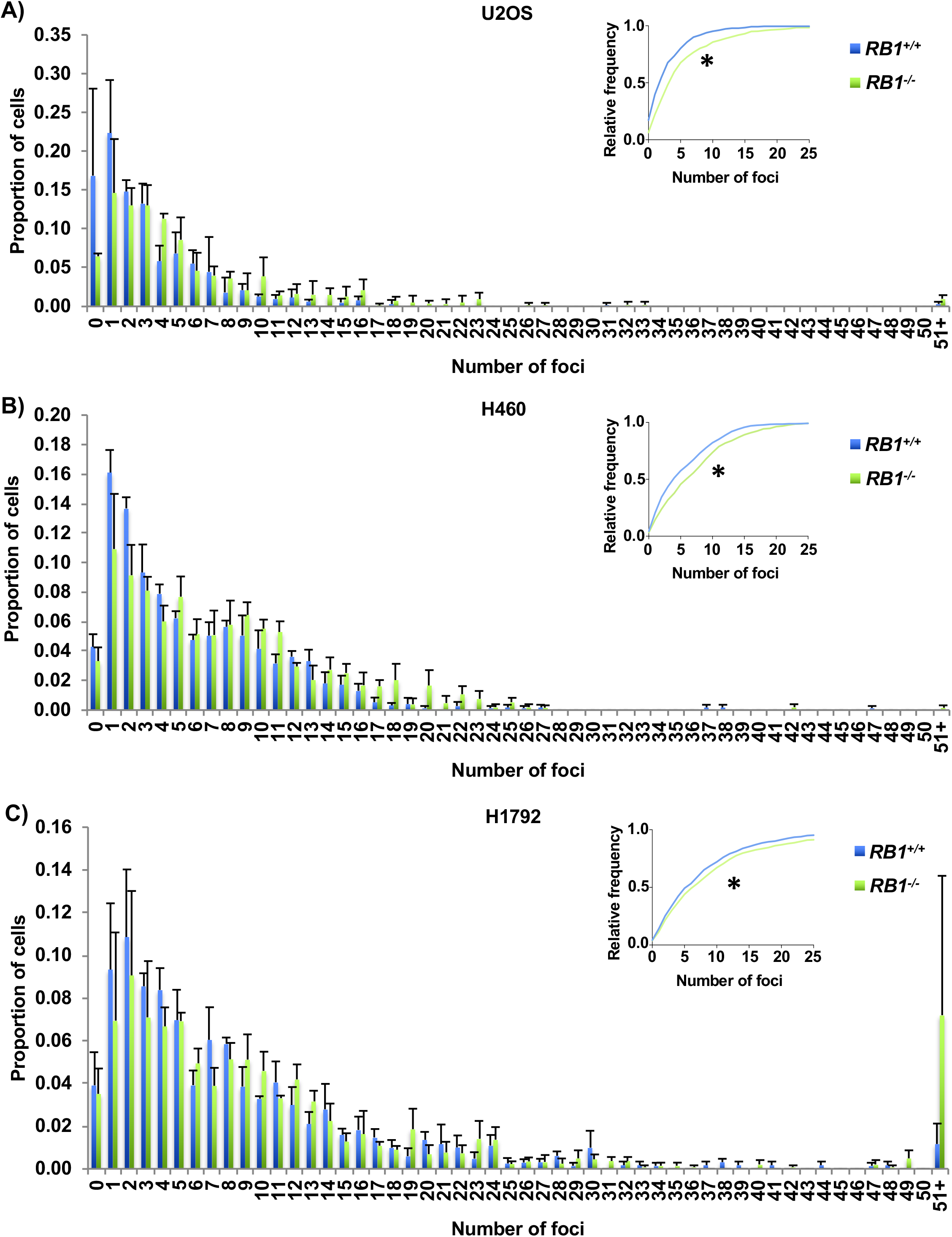
Spontaneous γH2AX foci are found in other induced *RB1* mutant cancer cell lines. **(A)** U2OS cells were transfected with CRISPR-Cas9 constructs targeting either a safe harbor site in the genome or the *RB1* gene. Three clones were selected for both control and knockout conditions and γH2AX foci were quantified by fluorescence microscopy. The average proportion of γH2AX foci for both *RB1* wild type and knockout genotypes are shown as histograms with standard error, while the cumulative relative frequency of foci is shown in the top right of the graph. Foci count distributions are significantly different as determined by Kolmogorov-Smirnov test. **(B)** Same experiment as in A, but using H460 cancer cells. **(C)** As in A, except with H1792 non-small cell lung cancer cells. All error bars are +1 SEM. *P < 0.05.

**Supplemental Figure 3.**
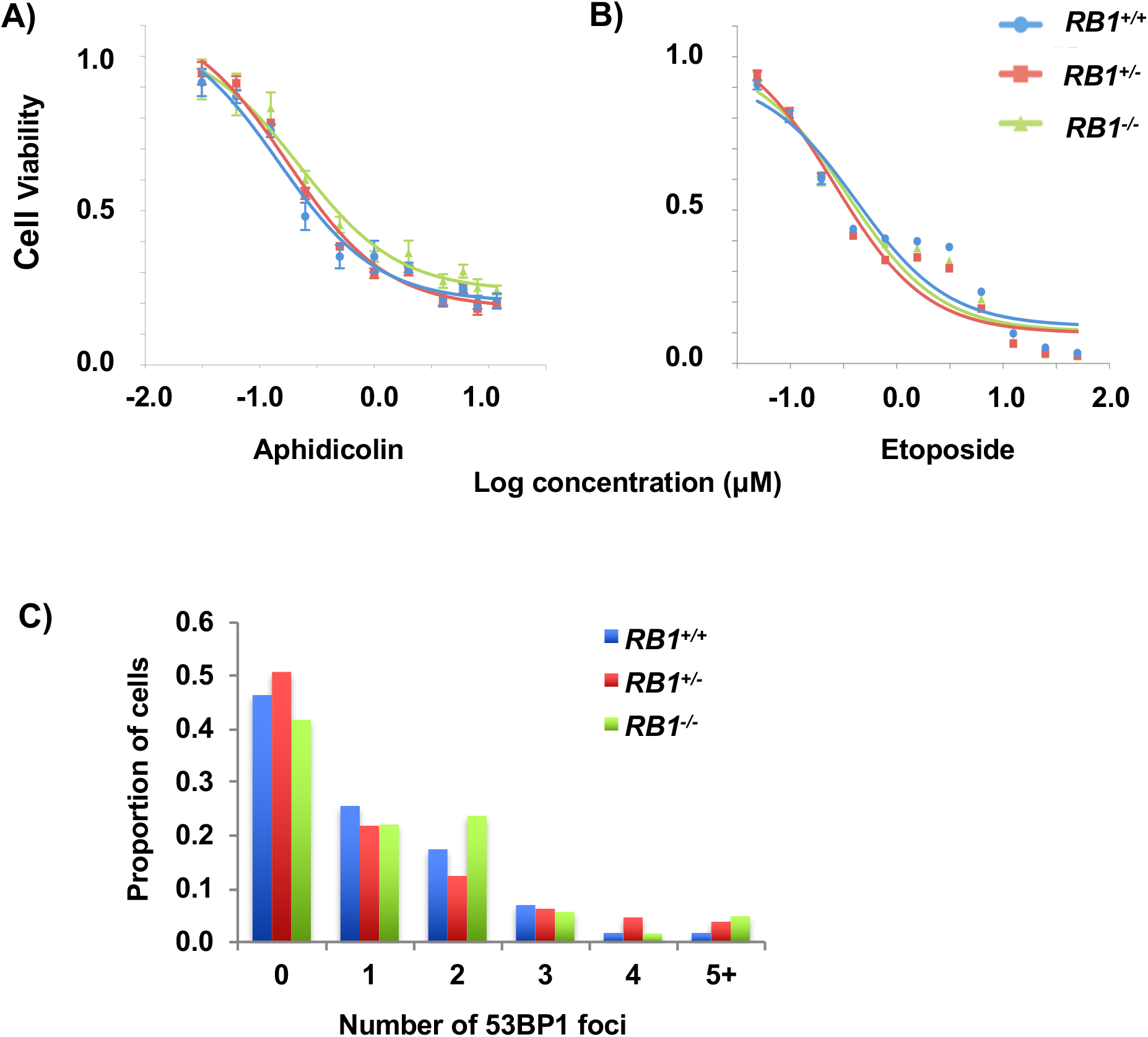
*RB1* mutant cells are not more sensitive to the direct induction of DNA breaks. **(A)** Aphidicolin and **(B)** etoposide were added to cultures of the indicated genotypes of cells. Viability was assessed after 72 h using alamarBlue and dose response curves were used to calculate IC50 values for each genotype. There are no differences between genotypes (as determined by one-way ANOVA). **(C)** 53BP1 foci were quantitated for each *RB1* genotype using the Focinator as with γH2AX. No significant differences were observed as determined by the Kolmogorov-Smirnov test. At least 120 cells from one U2OS clone were analyzed for each *RB1* genotype. All error bars are ±1 SEM.

**Supplemental Figure 4.**
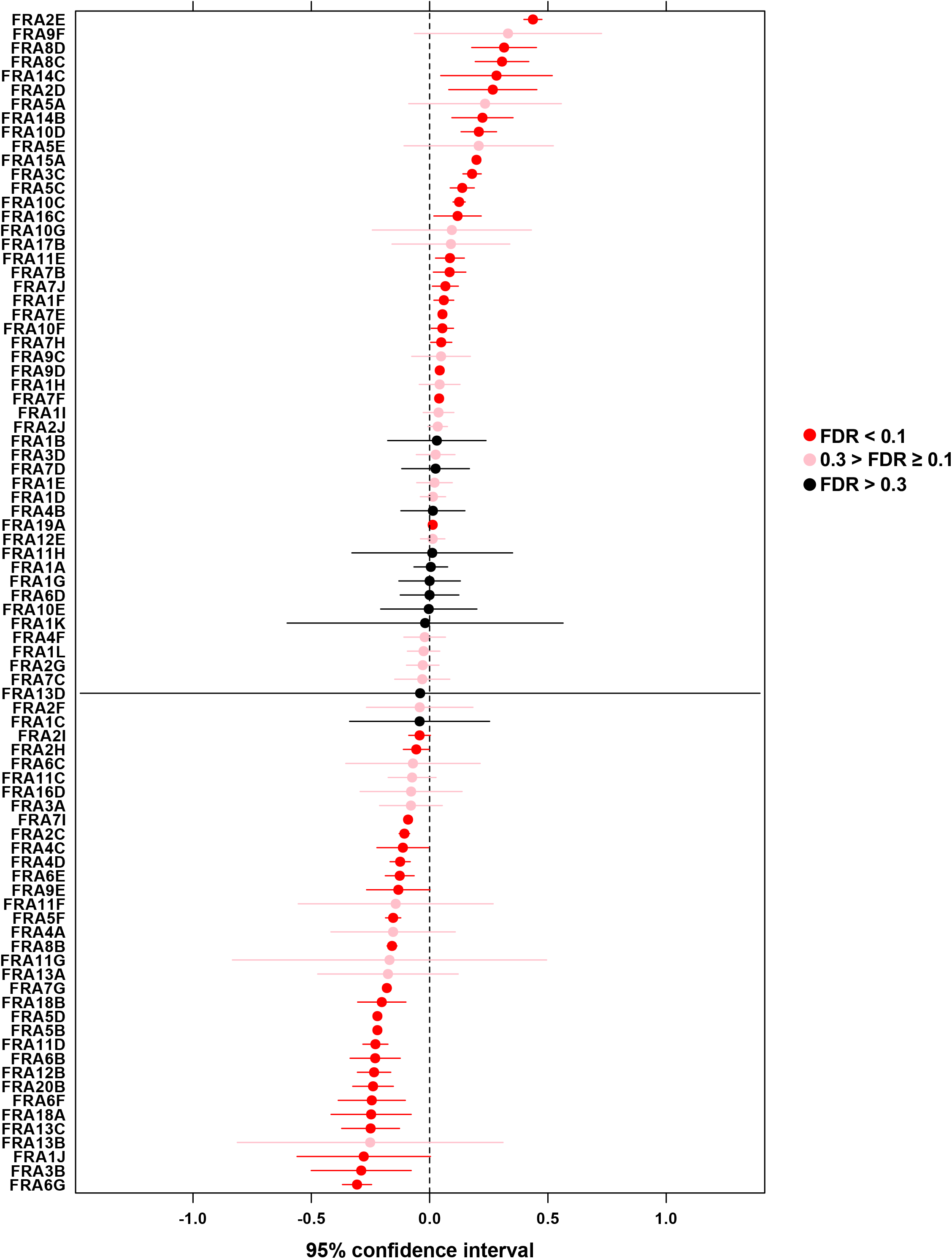
γH2AX ChIP-Seq read proportions in *RB1* mutants at CFS compared to wild type. Aligned γH2AX ChIP-Seq read proportions within common fragile sites (CFS) were first normalized to their respective inputs, and then *RB1*^+/−^ and *RB1*^−/−^ proportions were normalized to wild type and log2 transformed. A two-tailed one-sample *t*-test was performed to determine if the normalized mean read count proportions of the *RB1* mutants at the various CFS is equal to the normalized read count proportion of the corresponding CFS in the wild type. The dots in the middle of each segment are the mean values of the normalized, log2 transformed *RB1* mutant proportions and data are color-coded according to their FDR multi-test corrected *P*-values.

**Supplemental Figure 5.**
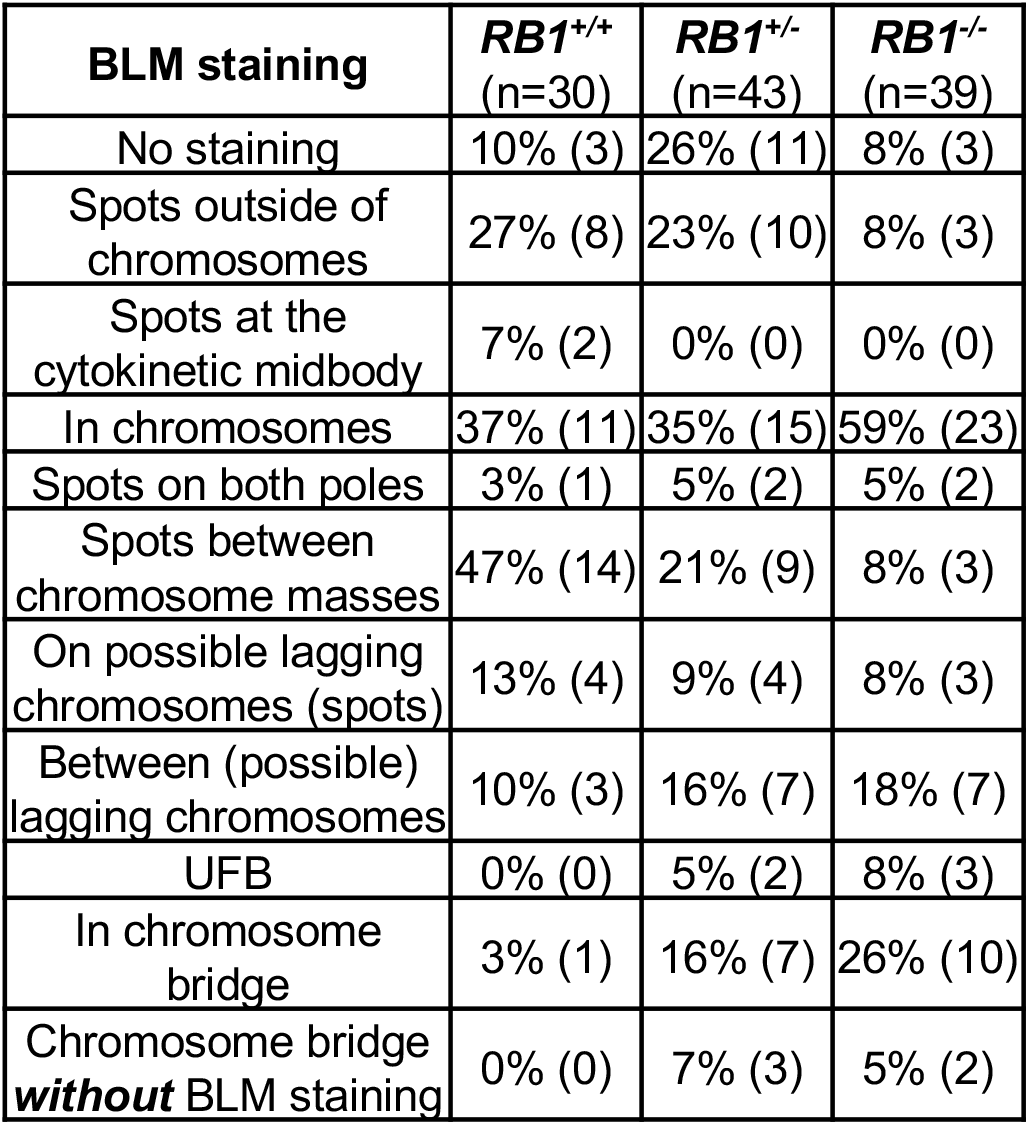
Quantitation of BLM staining patterns in anaphase. Cells in anaphase were imaged by fluorescence microscopy using DAPI and BLM from each *RB1* genotype, as in Figure 4. Images, as depicted in Figure 4C, were analyzed to determine the locations of BLM staining during anaphase. The categories are overlapping as all BLM staining events in the images were noted. The percent of images containing each type of staining are listed, with the number of images noted in brackets.

**Supplemental Figure 6.**
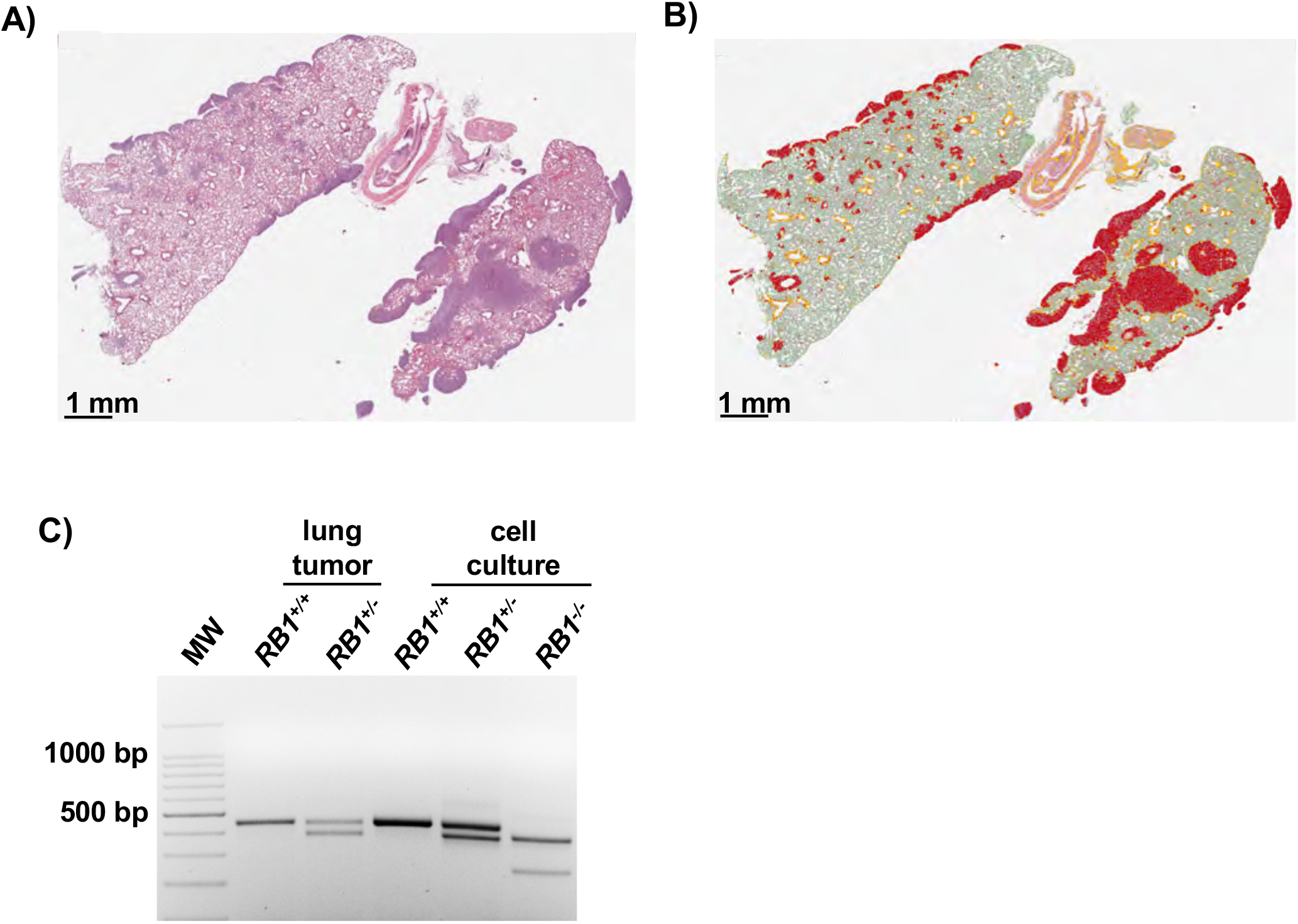
Confirmation of *RB1+* genotype in lung histology. **(A)** H&E staining of lung tissue with the highest number of *RB1*^+/−^ seeding events. **(B)** QuPath coloring to denote tumor tissue within these lungs. **(C)** DNA was extracted from clusters of tumor cells in paraffin embedded tissue containing these *RB1*^+/−^ cells, or a *RB1*^+/+^ control. PCR was performed to amplify exon 22 from recovered DNA and controls isolated from cell culture to verify the genotype of this sample.

